# Hepatitis B virus polymerase restricts LINE-1 retrotransposition

**DOI:** 10.1101/2021.05.07.443105

**Authors:** Yasuo Ariumi

## Abstract

Long interspersed element-1 (LINE-1, L1) retrotransposon composes about 17% of the human genome. However, genetic and biochemical interactions between L1 and hepatitis B virus (HBV) remain poorly understood. In this study, we found that HBV restricts L1 mobility without inhibiting the L1 promoter activity. Notably, HBV polymerase (Pol) strongly inhibited L1 retrotransposition in a reverse transcriptase (RT)-independent manner. Indeed, the ribonuclease H (RNase H) domain was essential for inhibition of L1 retrotransposition. L1 ORF1p RNA-binding protein predominantly localized into cytoplasmic RNA granule termed P-body. However, HBV Pol sequestered L1 ORF1p from P-body and colocalized with L1 ORF1p in cytoplasm, when both proteins were co-expressed. Altogether, HBV Pol seems to restrict L1 mobility through a sequestration of L1 ORF1p from P-body. Thus, these results suggest a novel function or activity of HBV Pol in regulation of L1 retrotransposition.

## 1. Introduction

Chronic infection with hepatitis B virus (HBV) is a major risk for hepatitis, cirrhosis, and the development of hepatocellular carcinoma (HCC). HBV infects more than 240 million people worldwide and remains a global public health problem (Rajoriya et al., 2017). HBV is a small enveloped DNA virus and belongs to the *Hepadnaviridae* family. The hepatocyte-specific bile acid transporter, sodium taurocholate co-transporting polypeptide (NTCP) serves as HBV receptor (Yan et al., 2012). The HBV genome (∼3.2kb) encodes four overlapping open reading frames (ORFs), C, P, S, and X, which express capsid-forming HBV core protein (HBc), DNA polymerase (Pol), small, medium and large HBV envelope proteins termed HBV surface antigens (HBs), L-HBs, M-HBs, and S-HBs, and regulatory HBV X protein (HBx), respectively (Seeger et al., 2015; Ko et al., 2017; Tsukuda and Watashi, 2020). Pol is the largest HBV protein, consisting of four functional domains; the terminal protein (TP) domain, which is essential for binding to HBV pregenomic RNA (pgRNA) and serves as a protein primer to initiate minus strand DNA synthesis; the spacer domain; the reverse transcriptase (RT) domain, which has both the RNA-directed DNA polymerase activity (reverse transcriptase, responsible for reverse transcription) and the DNA-directed DNA polymerase activity for viral replication; the ribonuclease H (RNase H) domain, which digests pgRNA after reverse transcription. The error prone HBV polymerase lacks proofreading activity causes the genetic variability.

Long interspersed element-1 (LINE-1, L1) is a mobile genetic element composing about 17% of the human genome (Ostertag et al., 2001). L1 encodes at least two open reading frames (ORFs), ORF1p with RNA binding domain and nucleic acid chaperone activity, and ORF2p with endonuclease and reverse transcriptase activities required for its retrotransposition (Mathias et al., 1991; Martin et al., 2001). L1 transcription occurs through promoter activity located in its 5’ untranslated region (UTR) (Swergold G.D., 1990). L1 mRNA assembles with ORF1p and ORF2p to form a ribonucleoprotein (RNP) complex in the cytoplasm (Doucet et al., 2010). Then, L1-RNP complex enters the nucleus in which integration occurs by a mechanism termed target-primed reverse transcription (TPRT). During TPRT, the L1 endonuclease creates a nicked DNA that serves as a primer for reverse transcription of L1 RNA, leading to integration of L1 cDNA into the human genome (Feng et al., 1996). The insertion of L1 into the human genome may cause genomic instability, genetic disorders, and cancers through insertional mutagenesis. Therefore, host cells have evolved several defense mechanisms protecting cells to restrict deleterious retrotransposition including epigenetic regulation through DNA or histone methylation and host restriction factors, such as apolipoprotein B mRNA editing enzyme catalytic polypeptide 3 (APOBEC3) family of cytidine deaminases, Moloney leukemia virus 10 (MOV10) RNA helicase, Rad18 DNA repair enzyme and sterile alpha motif and HD-containing protein 1 (SAMHD1) (Ariumi Y, 2016; Ariumi et al. 2018).

Human immunodeficiency virus type 1 (HIV-1) reverse transcriptase synthesizes a DNA copy of the HIV-1 genomic RNA and integrates into the human genome. Therefore, both HIV-1 and L1 might mutually influence their mobility in the human genome. In fact, we and other groups recently reported that HIV-1 affects L1 retrotransposition and expression (Jones et al., 2013; Iijima et al., 2013; Kawano et al., 2018). Similarly, HBV also has reverse transcriptase activity and synthesizes a DNA copy of HBV and integrates into the human genome. However, interaction between HBV and L1 are not well understood. Therefore, we investigated a cross talk of HBV with L1 in this study.

## 2. Material and methods

### 2.1. Cell culture

293T cells were cultured in Dulbecco’s modified Eagle’s medium (DMEM; Life Technology, Carlsbad, CA, USA) with high glucose supplemented with 10% fetal bovine serum (FBS) and 100 U/ml penicillin/streptomycin.

### 2.2. Plasmid construction

The payw1.2 (Scaglioni et al., 1997) (kind gift from Robert J. Schneider, New York University), pCMVayw1.2 HBV (kind gift from Stefan Wieland and Francis V. Chisari, Scripps Research Institute), pCMVayw1.2 YMHD (kind gift from Priscilla Turelli and Didier Trono, EPFL, Switzerland) were used (Biermer et al., 2003; Turelli et al., 2004). To construct pcDNA3-FLAG-HBc, a DNA fragment encoding HBc was amplified from payw1.2 WT (Scaglioni et al., 1997) by PCR using KOD-Plus DNA polymerase (TOYOBO, Osaka, Japan) and the following pair of primers: 5’-CGGGATCCAAGATGGACATCGACCCTTATA-3’ (Forward), 5’-CCGGCGGCCGCCTAACATTGAGATTCCCGA-3’ (Reverse). The obtained DNA fragment was subcloned into either the *Bam*HI-*Not*I sites of the pcDNA3-FLAG vector and the nucleotide sequences were determined by Sanger sequencing. To construct pcDNA3-HBx-FLAG or pcDNA3-HBV Pol-FLAG, a DNA fragment encoding HBx or HBV Pol was amplified from payw1.2 WT by PCR using KOD-Plus DNA polymerase and the following pairs of primers: HBx, 5’-CGAAGCTTAAGATGGCTGCTAGGCTGTGCT-3’ (Forward), 5’-CCGGCGGCCGCTTACTTATCGTCGTCATCCTTGTAATCGGCAGAGGTGAAAAAG-3’ (Reverse); HBV Pol, 5’-CGAAGCTTAAGATGCCCCTATCCTATCAAC-3’ (Forward), 5’-CCGGCGGCCGCTCACTTATCGTCGTCATCCTTGTAATCCGGTGGTCTCCATGCG-3’ (Reverse). The obtained DNA fragments were subcloned into either the *Hin*dIII-*Not*I sites of the pcDNA3 vector (Invitrogen). To construct several HBV Pol deletion mutants, pcDNA3-ΔRH-FLAG, pcDNA3-RH-FLAG, pcDNA3-RT-FLAG, pcDNA3-TP-FLAG or pcDNA3-ΔTP-FLAG, a DNA fragment encoding HBV Pol deletion mutant was amplified from payw1.2 WT by PCR using KOD-Plus DNA polymerase and the following pairs of primers: ΔRH, 5’-CGAAGCTTAACATGCCCCTATCCTATCAAC-3’ (Forward), 5’-CCGGCGGCCGCTCACTTATCGTCGTCATCCTTGTAATCCCGTTGCCGGGCAACGGG -3’ (Reverse); RH, 5’-CGAAGCTTACCATGCCAGGTCTGTGCCAAG-3’ (Forward), 5’-CCGGCGGCCGCTCACTTATCGTCGTCATCCTTGTAATCCGGTGGTCTCCATGCG-3’ (Reverse); RT, 5’-CGAAGCTTACCATGGAGGATTGGGGACCCT-3’ (Forward), 5’-CCG GCGGCCGCTCACTTATCGTCGTCATCCTTGTAATCCCGTTGCCGGGCAACGGG-3’ (Reverse); TP, 5’-CGAAGCTTAACATGCCCCTATCCTATCAAC-3’ (Forward), 5’-CCGGCGGCCGCTCACTTATCGTCGTCATCCTTGTAATCATCTTGTTCCCAAGAATA-3’ (Reverse); ΔTP, 5’-CGAAGCTTACCATGGAGGATTGGGGACCCT-3’ (Forward), 5’-CCGGCGGCCGCTCACTTATCGTCGTCATCCTTGTAATCCGGTGGTCTCCATGCG-3’ (Reverse). The obtained DNA fragments were subcloned into either the *Hin*dIII-*Not*I sites of the pcDNA3 vector (Invitrogen).

### 2.3. Immunofluorescence and confocal microscopy analysis

293T cells were grown on Lab-Tek 2 well chamber slide (Nunc, Thermo) at 2×10^4^ cells per well and transfected the next days using TransIT-LT1 transfection reagent. Two days after transfection, the cells were fixed in 3.6% formaldehyde in 1X phosphate-buffered saline (PBS), permeabilized in 0.1% NP-40 in 1X PBS at room temperature, and incubated with anti-HA antibody (3F10; Roche Diagnostics, Mannheim, Germany or HA-7; Sigma, Saint Louis, MI, USA), anti-DDX6 (A300-460A; Bethyl Lab, Inc., Montgomery, TX, USA) and/or anti-FLAG antibody (M2, Sigma) at a 1:300 dilution in 1X PBS containing 3% bovine serum albumin (BSA) at 37°C for 30 min. Cells were then stained with Donkey anti-mouse IgG (H+L) secondary antibody, Alexa Fluor 594 conjugate and Donkey anti-rat or anti-rabbit IgG (H+L) secondary antibody, Alexa Fluor 488 conjugate (Thermo Fisher Scientific Inc., Waltham, MA, USA) at a 1:300 dilution in 1X PBS containing 3% BSA at 37°C for 30 min. Nuclei were stained with DAPI (4’, 6’-diamidino-2-phenylindole). Following washing 3 times in 1X PBS, the coverslides were mounted on slides using SlowFade Gold antifade reagent (Life Technology). Samples were analyzed under a confocal laser-scanning microscope (FV1200; Olympus, Tokyo, Japan).

### 2.4. L1 retrotransposition assays

The luciferase-based retrotransposition assay was performed using the firefly luciferase-based retrotransposition reporter pYX014 or pYX017 plasmid (Xi et al., 2011). 293T cells were seeded in 24 well plates at 2×10^4^ cells per plate and then transfected the next day with pYX014 or pYX017 (100 ng) using TransIT-LT1 transfection reagent (Mirus Bio LLC, Madison, WI53711, USA). All transfections were performed using equal number of tested plasmid DNA molecules, with the addition of empty vector into the transfection mixtures to compensate for plasmid size differences and reach equal amounts of DNA quantities per condition. Results were obtained through three independent transfections. Luciferase assays were performed 72h after transfection using luciferase assay reagent according to the manufacturer’s instructions (Promega, Madison, WI, USA). A Lumat LB9507 luminometer (Berthold, Bad Wildbad, Germany) was used to measure firefly luciferase activity.

## 3. Results

### 3.1. HBV restricts L1 retrotransposition

To investigate whether HBV affects L1 retrotransposition activity, HBV molecular clone containing 1.2-fold over-length HBVayw genome, pCMVayw1.2 HBV WT (ayw strain) or pCMVayw1.2 YMHD, reverse transcriptase (RT)-inactive mutant (Biermer et al., 2003; Turelli et al., 2004) were co-transfected with pYX014 plasmid, which has the firefly luciferase-based retrotransposition reporter cassette (Xi et al., 2011) (Fig. 1A) into 293T cells. pCMVayw1.2 expresses the pregenomic (pg) RNA under control of a CMV promoter. pYX014 harbors the L1_RP_ 5’untranslated region (UTR) promoter as well as the firefly luciferase gene inserted in the 3’UTR of L1 in the opposite direction of L1 coding sequence (Xi et al., 2011). The luciferase activity could be detected only after L1 retrotransposition has happened. Notably, HBV could strongly inhibit the luciferase-based L1 retrotransposition activity in a RT-independent manner (Fig. 1A). Substitution of the L1_RP_ 5’UTR promoter (pYX014) by a strong CAG promoter resulted in an increase of the retrotransposition activity (pYX017) (Xi et al., 2011) (Fig. 1B). As well, HBV inhibited the retrotransposition activity (pYX017) (Fig. 1B), suggesting that this inhibitory effect on L1 retrotransposition is independent of the L1 5’UTR promoter activity. To determine which HBV protein inhibits L1 retrotransposition, HBx, HBV Pol, or HBc-expressing plasmid was co-transfected with pYX014. Notably, HBV Pol strongly inhibited L1 retrotransposition (Fig. 1C).

**Fig.1.**
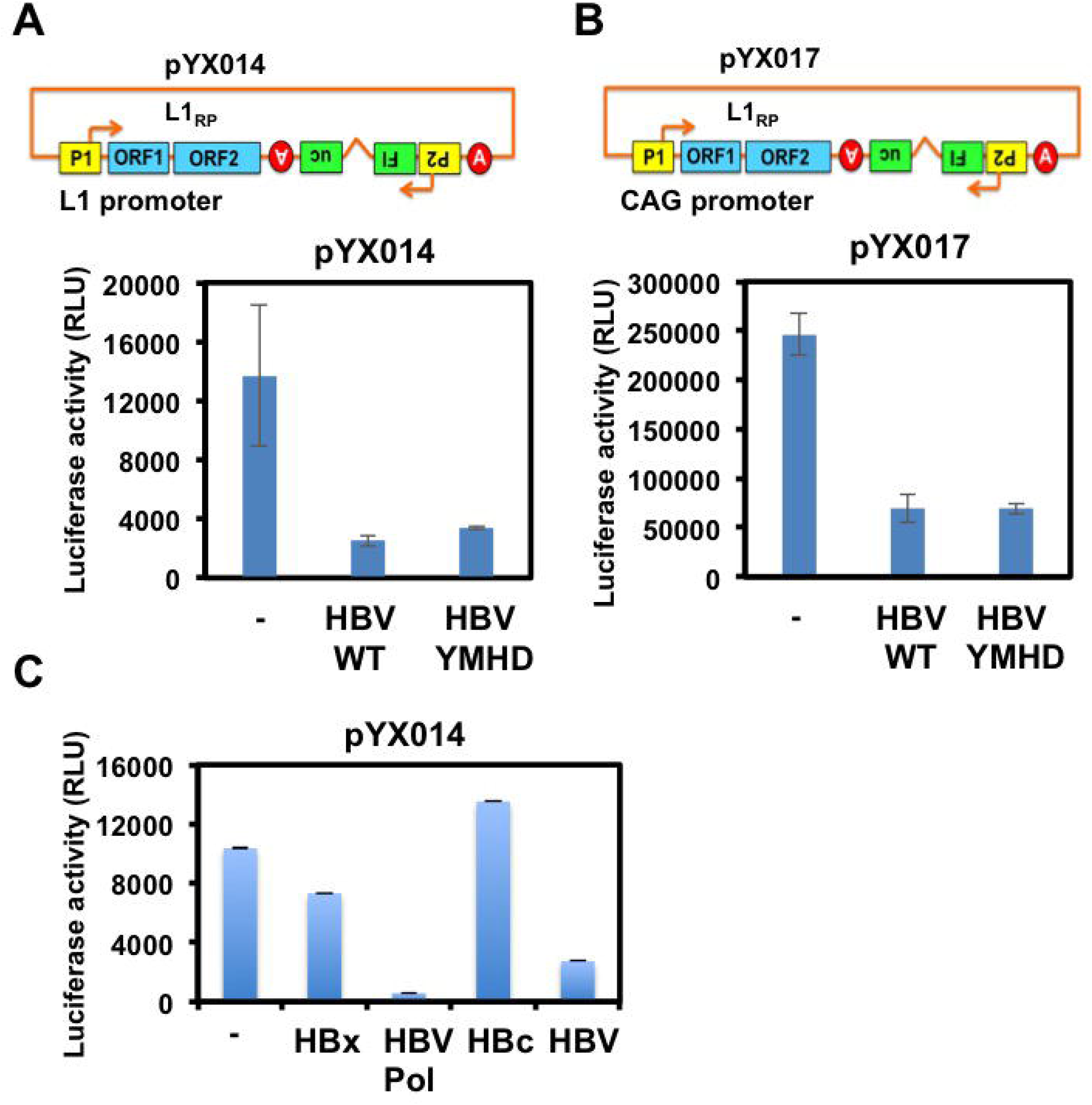
HBV Pol inhibits L1 retrotransposition. (A, B) Schematic representation of the firefly luciferase-based retrotransposition reporter cassette pYX014 and pYX017 (Xi et al., 2011). The L1_RP_ 5’UTR (pYX014) promoter was replaced by a strong CAG promoter and generated pYX017. 293T cells (2×10^4^ cells/well) were co-transfected with HBV molecular clone pCMVayw1.2 HBV WT or pCMVayw1.2 HBV YMHD (100 ng) and either pYX014 or pYX017 (100 ng) in triplicate. Three days after transfection, luciferase assays were performed. (C) 293T cells (2×10^4^ cells/well) were co-transfected with pcDNA3-HBx-FLAG, pcDNA3-HBV Pol-FLAG, pcDNA3-FLAG-HBc, or pCMVayw1.2 HBV WT (100 ng) and pYX014 (100 ng) in triplicate. Three days after transfection, luciferase assays were performed. All graphs show the mean (±SEM) firefly luciferase activity.

### 3.2. The RH domain of HBV Pol is essential for inhibition of L1 retrotransposition

Since HBV Pol consists four functional domains; the terminal protein (TP) domain, the spacer domain, the reverse transcriptase (RT) domain, and RNase H (RH) domain, we examined which domain is essential for inhibition of L1 retrotransposition. For this, we constructed several deletion mutants of HBV Pol (ΔRH, RH, RT, TP, ΔTP) (Fig. 2A). Each HBV Pol deletion mutant-expressing plasmid and pYX014 were co-transfected into 293T cells and examined their luciferase assays 72 h post-transfection. Consequently, the ΔTP mutant containing both RT and RH domains, and the RH domain alone strongly inhibited L1 retrotransposition like wild-type HBV Pol (Fig. 2B). In contrast, the RT or TP domain has no inhibitory effect (Fig. 2B). Thus, the RH domain of HBV Pol is essential for inhibition of L1 retrotransposition.

**Fig.2.**
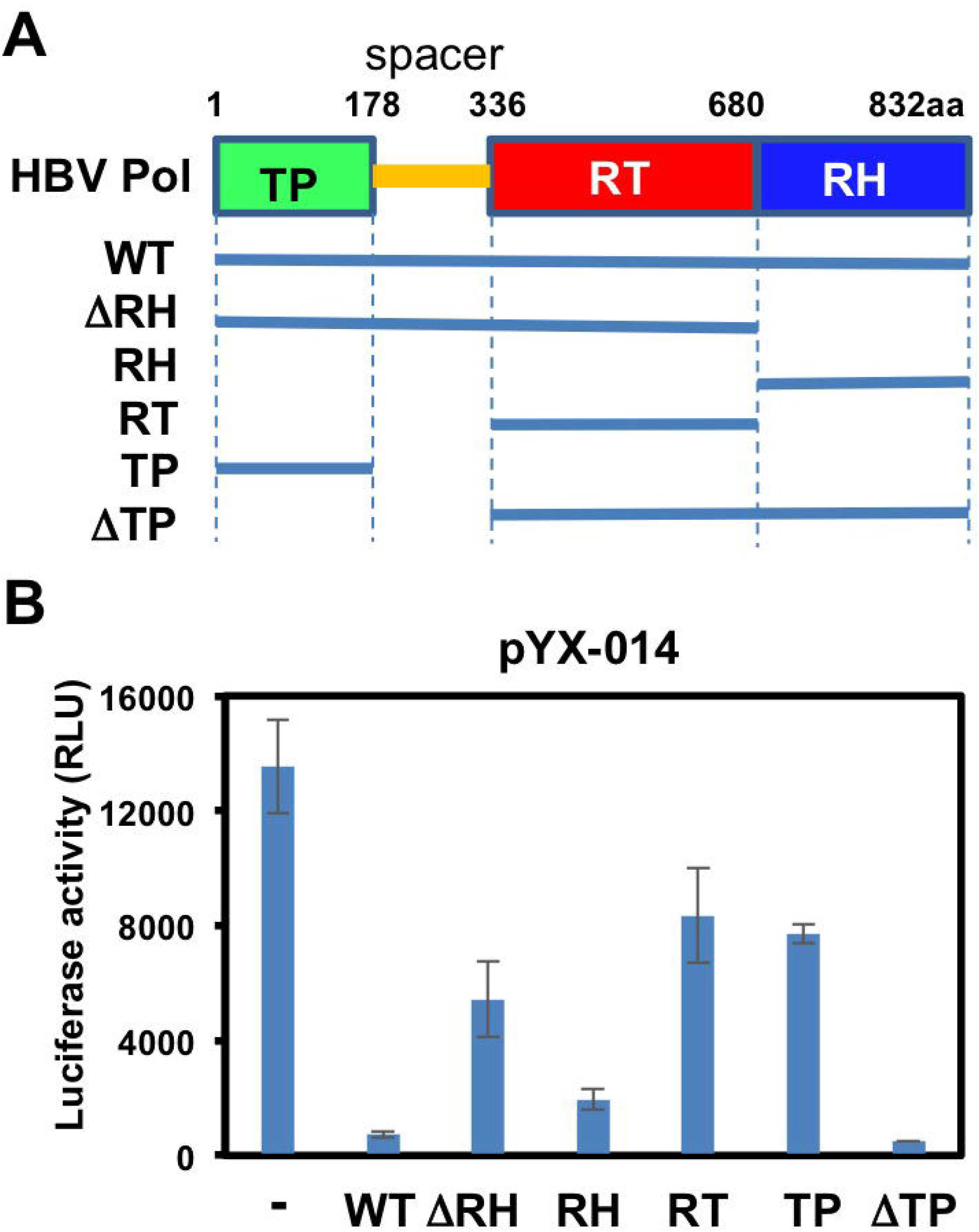
The RNase H domain is essential for inhibition of L1 retrotransposition. (A) Schematic representation of HBV Pol and the deletion mutants used in the present study. The terminal protein (TP), spacer, reverse transcriptase (RT), and RNase H (RH) domain are indicated. (B) 293T cells (2×10^4^ cells/well) were co-transfected with 100 ng of FLAG-tagged HBV Pol (WT) and the deletion mutant (ΔRH, RH, RT, TP, or ΔTP)-expressing plasmids and 100 ng of pYX014. Luciferase assays were performed three days after transfection in three independent experiments. Graph shows the mean (±SEM) firefly luciferase activity.

### 3.3. HBV Pol sequesters and colocalizes with L1 ORF1p

We next examined their subcellular localization of L1 ORF1p and HBV Pol or HBx by using a confocal laser scanning microscopy (FV1200, Olympus), when FLAG-tagged HBx or HBV Pol and HA-tagged L1 ORF1p were co-expressed. When L1 ORF1p alone was expressed in 293T cells, we observed that L1 ORF1p colocalizes with endogenous DDX6 RNA helicase (Fig. 3A). DDX6 predominantly localizes in P-body (Ariumi et al., 2011; Ariumi et al., 2018). Furthermore, L1 ORF1p also colocalized with FLAG-tagged AGO2, A3G-GFP, or FLAG-tagged MOV10 RNA helicase, well-known P-body components, indicating that L1 ORF1p localizes in cytoplasmic RNA granules termed P-bodies (Fig. 3A). Notably, HBV Pol sequestered L1 ORF1p from P-body and co-localized with L1 ORF1p in cytoplasm, when both proteins were co-expressed (Fig. 3B). However, HBV Pol did not sequester and colocalize with A3G-GFP, another P-body component (Fig. 3B). In contrast, HBx predominantly localized in cytoplasmic foci and did not co-localized with L1 ORF1p as well as A3G-GFP (Fig. 3B). Therefore, HBV Pol may inhibit L1 retrotransposition through a sequestration of L1 ORF1p from P-body.

**Fig.3.**
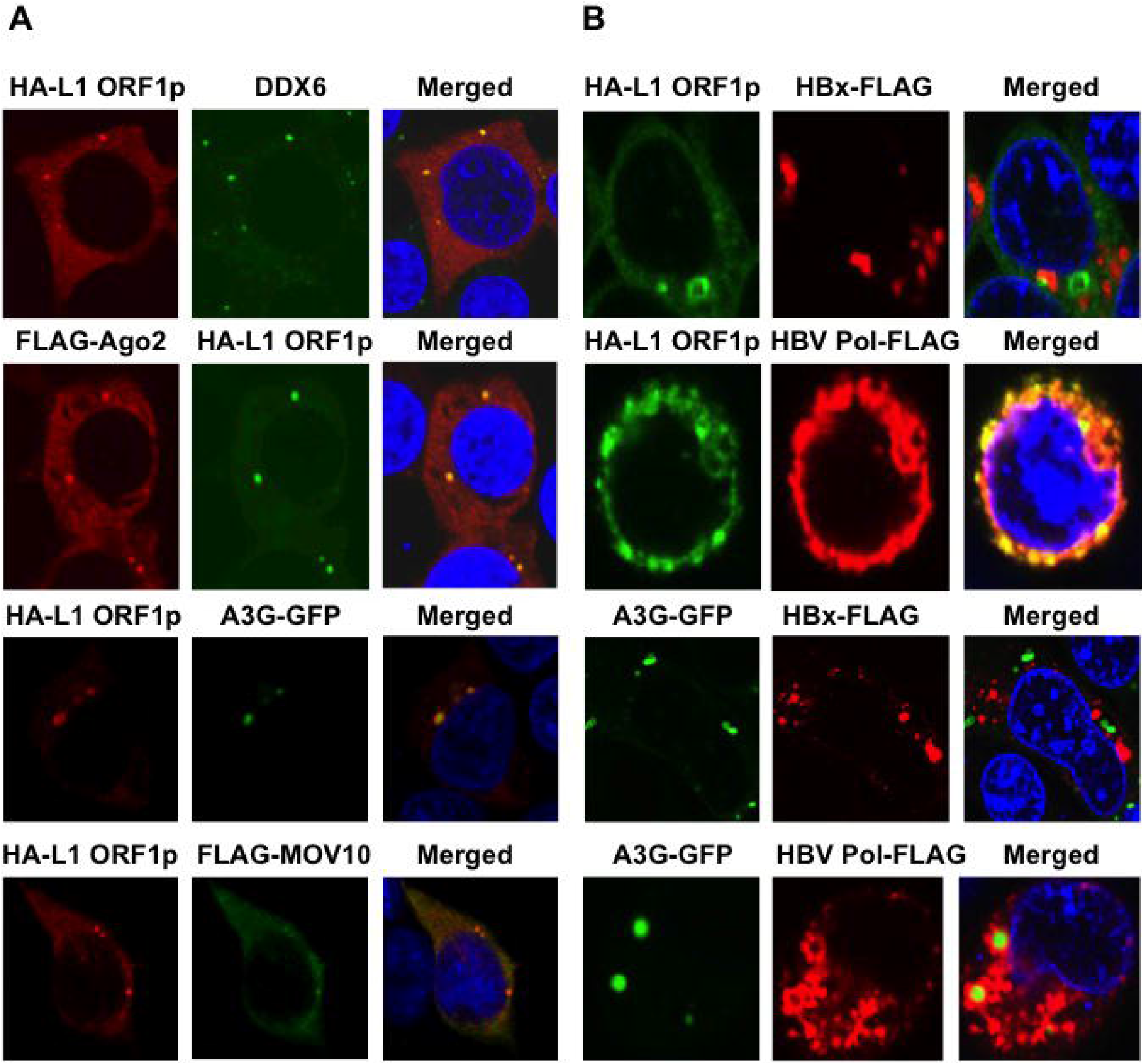
HBV Pol sequesters L1 ORF1p from P-body. (A) Subcellular localization of L1 ORF1p in P-body. 293T cells (2×10^4^ cells/well) transfected with 100 ng of pFLAG-Ago2, pA3G-GFP, pcDNA3-FLAG-MOV10 and/or 100 ng of pcDNA3-HA-L1 ORF1 (Ariumi et al. 2018; Kawano et al., 2018) were examined by confocal laser scanning microscopy (FV1200, Olympus). Cells were stained with anti-FLAG, anti-DDX6, and/or anti-HA antibodies and then visualized with Alexa Fluor 594 or Alexa Fluor 488. Nuclei were stained with DAPI. (B) Subcellular localization of HBx or HBV Pol and L1 ORF1p. 293T cells (2×10^4^ cells/well) cotransfected with 100 ng of pcDNA3-HBx-FLAG or pcDNA3-HBV Pol-FLAG and 100 ng of pcDNA3-HA-L1 ORF1 (Ariumi et al. 2018; Kawano et al., 2018) or pA3G-GFP. Cells were stained with anti-HA and anti-FLAG antibodies and then visualized with Alexa Fluor 594 or Alexa Fluor 488. Nuclei were stained with DAPI.

## 4. Discussion

In this study, I have, for the first time, demonstrated that HBV restricts L1 mobility. Notably, HBV Pol inhibited L1 retrotransposition in a RT-independent manner (Fig. 1). HBV Pol contains four functional domains; the terminal protein (TP) domain, the spacer domain, the reverse transcriptase (RT) domain, and RNase H (RH) domain. Indeed, the RH domain was essential for inhibition of L1 retrotransposition (Fig. 2). Therefore, HBV Pol may have a novel function or activity of HBV Pol in regulation of L1 retrotransposition.

L1 harbors its 5’UTR promoter to regulate L1 transcription (Swergold G.D. 1999). Several transcription factors including p53, RUNX3, SOX11, and YY1 positively regulate the L1 transcription (Athanikar et al., 2004; Ariumi Y, 2006; Becker et al., 1993; Harri et al., 2009; Yang et al., 2003; Tchenio et al., 2000). On the other hand, SOX2 and SRY negatively regulate the L1 transcription (Ariumi Y, 2006; Muotri et al., 2005; Tchenio et al., 2000). In this regard, HBV repressed L1 retrotransposition without inhibiting the L1 5’UTR promoter activity (Fig.1A and B), suggesting that this inhibitory effect on L1 retrotransposition is independent of the L1 5’UTR promoter activity. Since HBV Pol sequestered and colocalized with L1 ORF1p (Fig. 3), HBV pol may inhibit the L1 retrotransposition through the sequestration of L1 ORF1p from P-body. Consistent with our finding, Schöbel et al recently reported that hepatitis C virus infection restricts L1 retrotransposition by sequestration of L1 ORF1p in HCV-induced stress granules (Schöbel et al., 2021).

Chronic infection with HBV is a major risk for the development of hepatocellular carcinoma (HCC) (Rajoriya et al., 2017). HBV integration into the human genome was found in most HBV-related HCC and HBV integration has been implicated in the development of HCC. Although previous study reported that HBV integration occurs randomly without preferred integration site (Matsubara and Tokino, 1990), a recent high-throughput sequencing-based approaches identified recurrent integration sites in HCC (Ding et al., 2012). Indeed, telomerase reverse transcriptase (TERT) gene was identified as the most common recurrent HBV integration target sites. TERT encodes telomerase catalytic subunit, which is an essential component for maintenance telomere length. TERT plays a role in cellular senescence as well as immortality of various cells, such as cancer cells. Aberrant expression of TERT is associated with tumor development. Furthermore, seven integrations were found in the mobile genetic elements including L1, ERV1 and Alu (Ding et al., 2012). Similarly, L1 was identified as an HBV integration site in HBV-positive HCC cell lines by a recent whole-transcriptome study (Lau et al., 2014). Insertion of the gene encoding HBx into L1 in the human genome caused an oncogenic HBx-L1 chimeric RNA transcript (Lau et al., 2014). The HBx-L1 RNA transcript was detected in 23.3% of HCC patients and gave cancer-promoting properties through activation of Wnt/ß-catenin signaling pathway (Lau et al., 2014), suggesting that HBx-L1 increases risk of HCC development.

Furthermore, endogenous L1-mediated retrotransposition was identified in germline and somatic cells of HCC patients using retrotransposon capture sequencing (RC-seq) (Shukla et al., 2013). The germline L1 insertion in the tumor suppressor mutated in colorectal cancers (MCC) was detected in 21.1% of HCC, resulting in the aberrant expression of MCC. MCC expresses in liver and regulates oncogenic Wnt/ß-catenin signaling pathway in HCC. Moreover, suppression of tumorigenicity 18 (ST18), a member of the MYT1 zinc-finger transcription factor family, was activated by a tumor-specific somatic L1 insertion (Shukla et al., 2013), indicting that endogenous retrotransposition activates oncogenic pathways in HCC. Thus, L1-mediated retrotransposition seems to be involved in development of HCC. Therefore, interaction between HBV and L1 may shed light on the novel mechanism(s) how HBV regulates L1 retrotransposition as well as development of HBV-related HCC.

So far, several chronic viruses have been restricted L1 retrotransposition, including HIV-1, HCV, and HBV (Kawano et al., 2018; Schöbel et al., 2021; this study). First, HIV Vpr inhibited L1 retrotransposition in a cell cycle dependent manner, since Vpr is known to regulate host cell cycle (Kawano et al., 2018). In addition, Vpr associated with L1 ORF2 reverse transcriptase and suppressed its RT activity, however, Vpr did not interact with L1 ORF1p (Kawano et al., 2018). On the other hand, both HBV and HCV are causative agents for HCC and chronic hepatitis. HCV and HBV restricted L1 retrotransposition through the sequestration of L1 ORF1p (Schöbel et al., 2021; this study). Thus, the interaction between endogenous transposable elements and exogenous viruses might contribute to disease progression.

## Conclusions

HBV, at least, restricts L1 mobility by three possible mechanisms; (1) HBV represses the L1 retrotransposition without inhibiting its promoter activity, (2) HBV Pol inhibits L1 retrotransposition using RNase H domain, (3) HBV Pol inhibits L1 retrotransposition through sequestration of L1 ORF1p from P-body. These results suggest a novel function or activity of HBV Pol in regulation of L1 retrotransposition.

## Acknowledgements

We thank Drs. Didier Trono, Priscilla Turelli, Wenfeng An, Haruhiko Siomi, Robert J. Schneider, Stefan Wieland, and Francis V. Chisari for reagents. This work was supported by the Japan Society for the Promotion of Science (JSPS) KAKENHI Grant Number JP18981606, by the Research Program on Hepatitis B Grant Number JP17929672 from Japan Agency for Medical Research and Development, AMED.

